# EncoMPASS: an Encyclopedia of Membrane Proteins Analyzed by Structure and Symmetry

**DOI:** 10.1101/391961

**Authors:** Antoniya A. Aleksandrova, Edoardo Sarti, Lucy R. Forrest

## Abstract

Protein structure determination and prediction, active site detection, and protein sequence alignment techniques all exploit information about protein structure and structural relationships. For membrane proteins, however, there is no agreement among available online tools for highlighting and mapping such structural similarities. Moreover, no available resource provides a systematic overview of quaternary and internal symmetries, and their orientation with respect to the membrane, despite the fact that these properties can provide key insights into membrane protein function. To address these issues, we created the <u>Enc</u>yclopedia <u>o</u>f <u>M</u>embrane <u>P</u>roteins <u>A</u>nalyzed by <u>S</u>tructure and Symmetry (EncoMPASS), a database for relating integral membrane proteins of known structure from the points of view of sequence, structure, and symmetry. EncoMPASS is accessible at https://encompass.ninds.nih.gov and its contents can be easily downloaded. This allows the user not only to focus on specific systems, but also to study general properties of the structure and evolution of membrane proteins.

**Highlights:**

-EncoMPASS relates and analyzes known structures of membrane proteins
-Structure and sequence similarity is assessed through alignments and topology considerations, not clustering
-Symmetry is detected based on CE-Symm and SymD using a multi-step procedure

## 1. Introduction

Integral membrane proteins constitute 20-30% of the genome of any given organism (Stevens and Arkin, 2000). Moreover, they are targeted by around half of all FDA-approved drugs (Cournia *et al.*, 2015) and physiologically-relevant small ligands (Bull and Doig, 2015), highlighting the significance of these proteins in both cell biology and pharmacology (Davey, 2004). The chemical and spatial characteristics of their environment neatly separate this class from the rest of the proteome: in the lipid bilayer, proteins acquire distinct functions, as well as a predisposition for intramolecular and quaternary symmetries that reflects the geometric constraints of the membrane (Bowie, 2001; Choi *et al.*, 2008; Myers-Turnbull *et al.*, 2014, Forrest 2015). A detailed understanding of these unique features relies on structural data, and yet the hydrophobicity of the lipid bilayer environment has also rendered membrane proteins historically elusive to structure determination. Fortunately, the number of reported membrane protein structures is now increasing at about the same pace as that of the entire proteome (blanco.biomol.uci.edu/mpstruc/, www.rcsb.org/stats/growth/overall). The resultant proliferation of data presents new challenges and opportunities for classification and analysis.

Several databases dedicated to structures of membrane proteins have been developed, all of which rely on predictions of the positioning of the protein in the membrane. Among the most well-known databases are PDBTM (Kozma *et al.*, 2013), OPM (Lomize *et al.*, 2012) and MemProtMD (Stansfeld *et al.*, 2015), which respectively use objective functions, free energy calculations, and coarse-grained molecular dynamics simulations, to predict the orientation of each protein in the lipid bilayer. None of these databases classify the structures or assign structural relationships between the proteins that they contain. Although the well-known protein classification schemes (SCOP (Hubbard *et al.*, 1997), CATH (Orengo *et al.*, 1997), DALI (Holm and Sander, 1995)) include classes for membrane proteins, in practice such whole-protein classifications have been found to be inconsistent (Neumann *et al.*, 2010); the fold assignment of domains with less than six transmembrane helices, which constitute the vast majority of all membrane-spanning domains (Frishman, 2010), is particularly problematic. More generally, doubts have been raised over the feasibility that a structural classification can be both rigidly-defined and accurate (Valas *et al.*, 2009; Bourne and Shindyalov, 2003), as it has become clearer that protein structures are distributed continuously over structure space (Skolnick *et al.*, 2009) and therefore cannot be readily organized into a set of discrete classes. As Petrey and Honig suggest, more dynamic approaches that integrate several different kinds of annotations, as well as residue-level estimators, are needed in order to increase database accuracy and robustness (Petrey and Honig, 2009).

Symmetry is another central feature of membrane proteins that available databases do not fully consider. Symmetry and pseudo-symmetry are observed not only within multisubunit assemblies, but also in the repetition of internal structural or sequence elements. Moreover, these features are often intimately associated with function (Goodsell and Olson, 2000; Forrest, 2015; Balaji, 2015). Although sophisticated algorithms exist that detect repeated elements in proteins (Kim *et al.*, 2010; Myers-Turnbull *et al.*, 2014), there has been limited effort to investigate their relevance to membrane proteins, especially with respect to the relationship of the symmetry to the membrane itself. In particular, there is no available database that systematically catalogues the symmetric features of this class of proteins, and that could be used as a foundation for functional and evolutionary studies of its members.

To address the aforementioned issues, we present here the Encyclopedia of Membrane Proteins Analyzed by Structure and Symmetry (EncoMPASS). EncoMPASS is a fully-automated database that provides a flexible representation of the structural relationships between membrane protein structures, combined with detailed information about their quaternary and internal symmetries. To ensure maximum coverage and accuracy, the EncoMPASS library combines data from the OPM, PDB, and PDBTM databases, including the predicted orientation of each protein in the lipid bilayer. The identification of structural relationships holds a number of challenges, not least the large number of false positives and false negatives that would result from attempting every possible structural comparison, and that could significantly compromise the accuracy of subsequent analysis. As a countermeasure, we select a subset of relatable membrane-spanning chains based on their transmembrane (TM) topologies. The structural alignments are then carried out by the program Fr-TM-Align (Pandit and Skolnick, 2008), which provides two measures of structural similarity, namely the template model score (TM-score, (Zhang and Skolnick, 2004)) and root mean squared deviation (RMSD). However, these scores may not appropriately identify relationships between protein structures of the same sequence that have undergone large conformational changes. We therefore also record the percentage of identical residues based on alignments of sequences obtained with MUSCLE (Edgar, 2004) for every pair of membrane protein structures.

The structural similarity comparisons result in a network of similarity scores between all structures, but do not provide information on the symmetries inherent to the structure. Thus, all structures are analyzed at the level of multi-subunit complexes, to identify intersubunit or “quaternary” symmetries, as well as of individual membrane-spanning chains, to identify intra-subunit or “internal” symmetries. However, the coverage and type of symmetry varies immensely among membrane protein structures, ranging from perfect symmetries to extreme asymmetries (Forrest, 2015), such that no single methodology is currently capable of recognizing all symmetries of interest. We thus decided to implement a multi-stage approach combining the strengths of two symmetry-detection algorithms, CE-Symm (Myers-Turnbull *et al.*, 2014; Bliven *et al.*, 2018) and SymD (Kim *et al.*, 2010). Additional analysis places the results in the context of their lipid bilayer environment, resulting in more functionally- and evolutionarily-meaningful descriptions of the symmetric relationships. Finally, by utilizing the aforementioned structural and sequence relationships, we identify symmetries that would otherwise remain undetected, particularly those symmetries that have become masked by conformational change.

In the following section, we briefly describe the key features of the EncoMPASS database.

## 2. Results

### 2.1. A uniform and consistent dataset of membrane proteins

The first aim of EncoMPASS is to curate a dataset of embedded membrane protein structures with accurate atomic coordinates and biological units that will then permit robust structure alignment and symmetry recognition analyses. We avoid structural uncertainty by selecting only X-ray structures and by discarding entries with resolution >3.5 Å or polypeptide chains with large missing segments (>100 consecutive amino acids). While both the OPM and the PDBTM databases provide sets of embedded membrane proteins, the two databases are not entirely consistent with each other in coverage or annotation and each suffers from its specific drawbacks (Shimizu et al., 2018). Hence, we developed an automated procedure that takes both databases into consideration when building our dataset. The OPM database was our primary source, since all its entries undergo both automatic processing and manual curation steps. Unfortunately, manual curation also renders the data prone to human error and therefore the procedure for generating the EncoMPASS dataset aims to miminize the impact of such errors. For example, to counteract errors in the biological unit we compare the OPM protein assembly with the first BIOMOLECULE record in the PDB file and, in the case of a mismatch, we trigger a sequence of additional checks, including an updated version of the biological unit selection procedure used in the PDBTM (Kozma et al., 2013) (see STAR Methods, Fig. 1).

**Figure 1.**
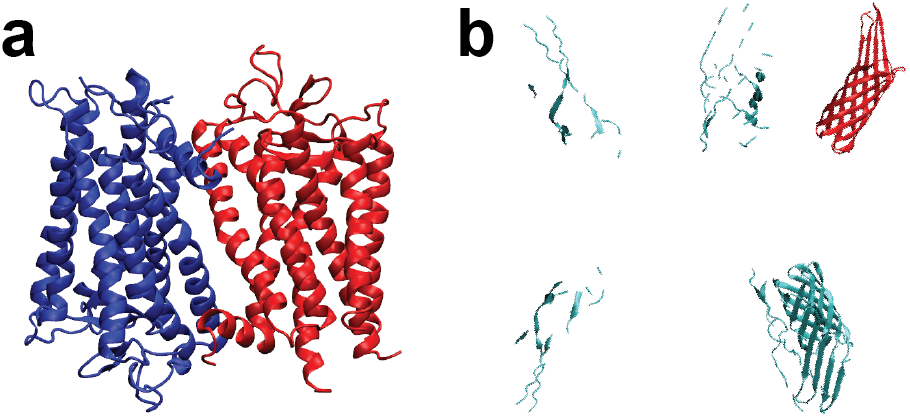
Examples of biological assembly errors. (a) An incorrect oligomeric state was indicated for bovine rhodopsin (PDB code 1F88). The assembly indicates an antiparallel dimer instead of a monomer or a parallel dimer (whose functional relevance is disputed). (b) An incorrect geometrical transformation matrix reported in the PDB file (PDB code 1QJ9) leads to an unfeasible assembly of the outer membrane protein OMPX from *E. coli.*

Since a complete list of erroneous biological assembly annotations is not available, we are unable to provide an absolute success rate for this procedure. Nevertheless, by visual inspection, we found 13 cases with trivial geometrical issues (STAR Methods). All these cases were identified and corrected by our procedure. In the final EncoMPASS database, out of 509 structures which were identified as potentially problematic, 7 were eventually considered correct, 442 were successfully modified, and 60 were discarded.

Once all the structures selected for checks in the previous stages have been either discarded or corrected, those that subsequently require positioning in the membrane are processed with PPM (Lomize *et al.*, 2012), the empirical free-energy minimization algorithm used in OPM.

The EncoMPASS database reports all and only the structures that have been successfully inserted in the PPM model lipid bilayer, either taken directly from the OPM database or inserted with the related PPM algorithm. As of May 15^th^ 2018, EncoMPASS contains 2344 complexes, containing a total of 7560 membrane-spanning single-chain subunits.

### 2.2. Transmembrane topologies and complex sizes

The EncoMPASS dataset contains structures with a wide range of complex sizes and number of membrane-spanning segments. The choice to base the design of EncoMPASS not only around each complex, but also on the component TM chains, enabled a detailed analysis of the number of segments in each chain and each complex, as well as the relationship between complex formation and number of TM regions (Fig. 2). For example, 33% of the complexes in the dataset contain only one subunit that crosses the membrane, and that subunit, in 80% of cases, also constitutes the only chain of the protein, i.e. monomers with a single membrane span. Examining individual protein chains, there is a prevalence for subunits that contain 1 or 2 TM regions (35% of the database), although these chains usually belong to larger complexes (Fig. 2b).

**Figure 2:**
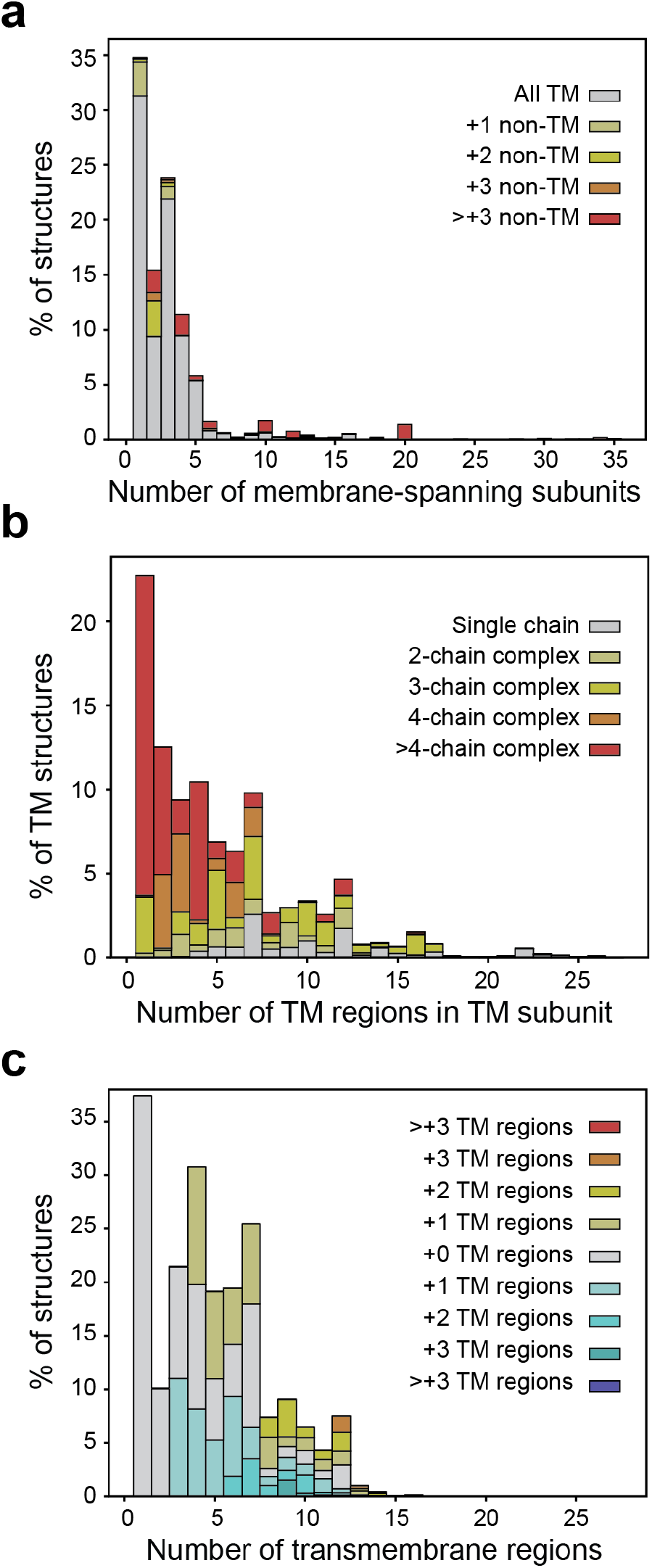
Relationships between the number of protein subunits that span the membrane, the number of TM regions in those subunits, and multi-subunit complex formation. (a) The fraction of chains or subunits in a membrane protein structure (both monomers and multi-subunit complexes) that are membrane-embedded. Coloring indicates the total number of subunits or chains in the complex to which each chain belongs. For example, the first column shows that among all the complexes with one of those chains in the membrane, the vast majority are monomers, followed by smaller percentages of dimers and trimers. (b) Division of all membrane-embedded subunits or chains according to the number of TM regions in each chain. Coloring indicates the total number of subunits or chains in the complex to which each chain belongs. For example, the first column shows that among protein chains in the database with 1 TM region, none belongs to a complex with <3 membrane-embedded subunits, while many more chains belong to complexes with >5 TM chains. (c) Nature of the comparisons between membrane protein structures following the selection criteria described in STAR methods. The y-axis indicates the fraction of all pairwise structural alignments that were performed between a-helical structures. The x-axis indicates the number of membrane-spanning regions in the protein chain of interest. Thus, fewer comparisons were carried out for structures with higher numbers of transmembrane helices, reflecting their distribution in the database. To identify whether each structure is compared to other structures with a similar number of TM helices, the color coding indicates the number of TM regions of the second TM chain in the pair.

### 2.3. Analysis of structural relationships

A major feature of EncoMPASS is the structural and sequence similarities among membrane TM chains. The simplest way to obtain this information would be to perform all pairwise structural alignments between any two single-chain subunits in the database and then assess the quality of each superposition. However, this simple brute-force approach would result in a large number of comparisons between structures that are plainly incompatible. In those cases, structural alignment algorithms are likely to find false positive similarities. To limit the number of compared structures, a straightforward strategy would be to compare chains only if they contain the same number of transmembrane (TM) regions. However, this strategy is problematic: first, such a methodology is not robust to errors in assigning the number of TM regions, even ignoring the unfortunate fact that there is no widely-accepted definition of a TM region. More importantly, even in the case of a perfectly accurate TM-region estimator, this simple strategy would disregard all similarities between chains in which single TM elements have been appended, inserted, or deleted during evolution.

A further complexity is that EncoMPASS contains an abundance of TM chains with very simple transmembrane topologies, i.e. with one or two membrane-spanning segments (Fig. 2c). Such protein chains are particularly prone to be inappropriately categorized with other structures, since there are many more solutions to match each secondary structure element to elements of another, more complex, structure. In particular, all pairs of 1-TM and 2-TM chains could potentially be successfully superposed, without indicating a true structural relationship. In such cases, the presence of domains outside of the membrane region can be used to assess whether it is reasonable to carry out a structural comparison. We thus restrict the number of allowed comparisons between 1-TM and 2-TM chains by considering their transmembrane topologies as well as the size of their extramembrane domains (see STAR Methods).

When comparing protein chains with >2 membrane-spanning regions, the potential for such false positives is smaller. However, more complex topologies are more prone to incorrect assignment of the number of TM regions. Moreover, it is not uncommon for trivially structurally-related proteins to have different numbers of TM regions. To account for this, we allow greater flexibility in the selection of chains for comparison if they contain >3 TM regions (see STAR Methods). Overall, this strategy reduces the number of comparisons between a-helical proteins from 49×10^6^ to 9×10^6^, while still allowing for pairs of structures with the same or similar numbers of TM segments to be compared (Fig. 3c).

**Figure 3:**
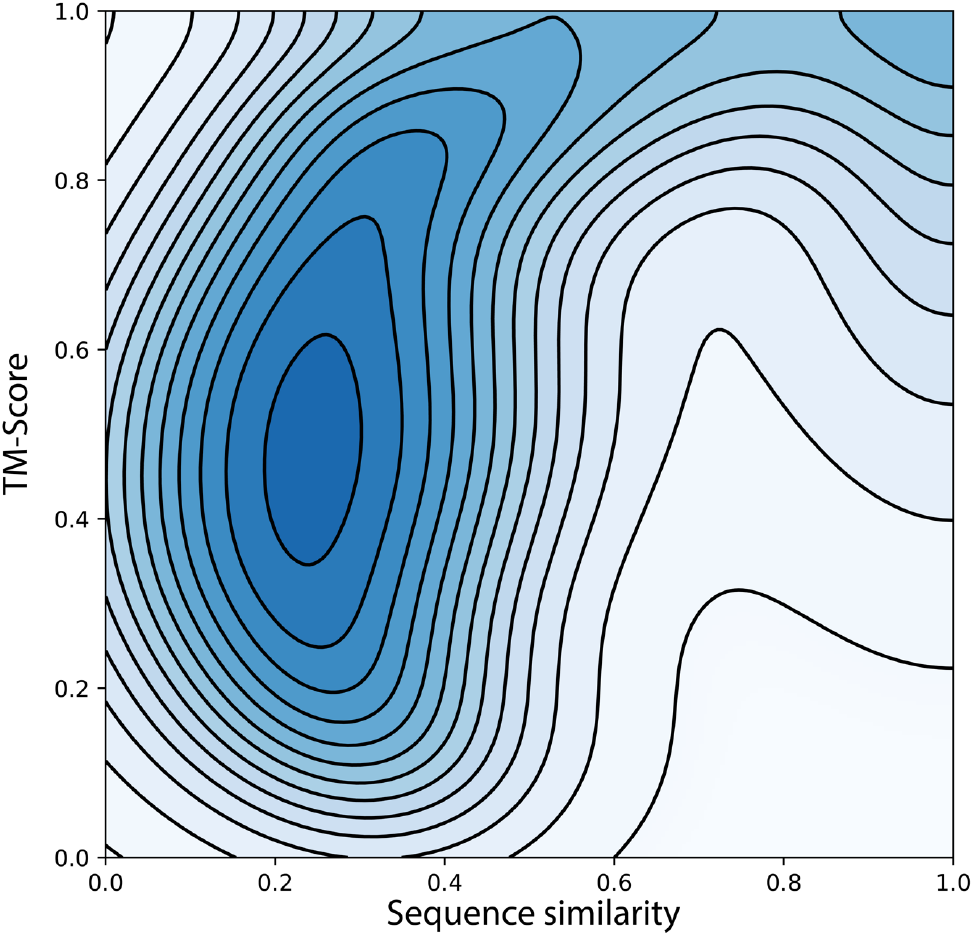
Structural and sequence similarity of all pairs in the EncoMPASS dataset. Contours indicate the probability density of the pairwise alignments having a given value of sequence identity and TM-score. The sequence identity is taken from the pair-wise MUSCLE alignment.

For membrane proteins with predominantly β-stranded folds, we carry out all-versus-all comparisons between their TM chains, since they are relatively scarce. However, comparisons between a-helical and β-stranded proteins are avoided. In total, we performed 9,419,278 structure and sequence alignments, which represent 16% of all possible pairwise alignments in the database.

Once all selected structure alignments have been carried out, we report the TM-score and RMSD of the alignments. We also report the sequence identity extracted from the structure-based sequence alignment provided by Fr-TM-Align. Separately, the sequences of the two chains are aligned with MUSCLE, i.e. without taking into account structural information, and the sequence identity is also calculated from this alignment. Analyzed over the entire database, it is clear that for most of the selected pairs, both sequence and structure similarity are low (Fig. 3). However, there is also a large population of pairs of chains that are dissimilar, as measured by sequence identity (<30%), and yet have structures that are considered similar (i.e. TM-score >0.6).

To organize this information for each chain in an intuitive manner, we provide three types of graph for each TM chain of interest. Each plot summarizes a different aspect of the structure and sequence relationships of that chain relative to the rest of the database. We first relate the sequence and structure similarity of all alignments carried out with the TM chain of interest (Fig. 4a) and, for reference, superpose those results onto the underlying density plot for the entire database (c.f. Fig. 3). As an example, the Kv1.2 potassium channel structure (PDB code 2A79 chain J) has several close sequence relatives (>90% identity) in the database that have the same number of TM segments, but quite different structures (TM-scores ~0.65), in addition to >1000 structures that are dissimilar both structurally and at the sequence level. Note that this representation provides no information about whether two points positioned close to each other originate from two similar structures or rather from two structures that just happen to be similarly related to the TM chain of interest. To examine the relationships between all related chains, we provide a polar representation (Fig. 4b), where the distance between every pair of points is proportional to the similarity of the corresponding structures. In this example, the structural relatives of Kv1.2 appear to cluster in 3 small groups, two of which are more similar to each other than the third, but all of which have the same number of TM segments. These plots provide overviews of the structure relationships, but it is not clear whether certain parts of the structure are more similar than others. To this end, we also provide a detailed residue-wise description of the structural similarity of the TM chain to all compared structures (Fig. 4c), where each line represents the Cα-Cα distance of a pair of aligned structures. Thus, in the case of the Kv1.2 structure, the closest structural relatives appear to be well-aligned in the N-terminal ~130 residues, while none of the available structures matches the C-terminal half of the first TM segment (residues ~350-280).

**Figure 4:**
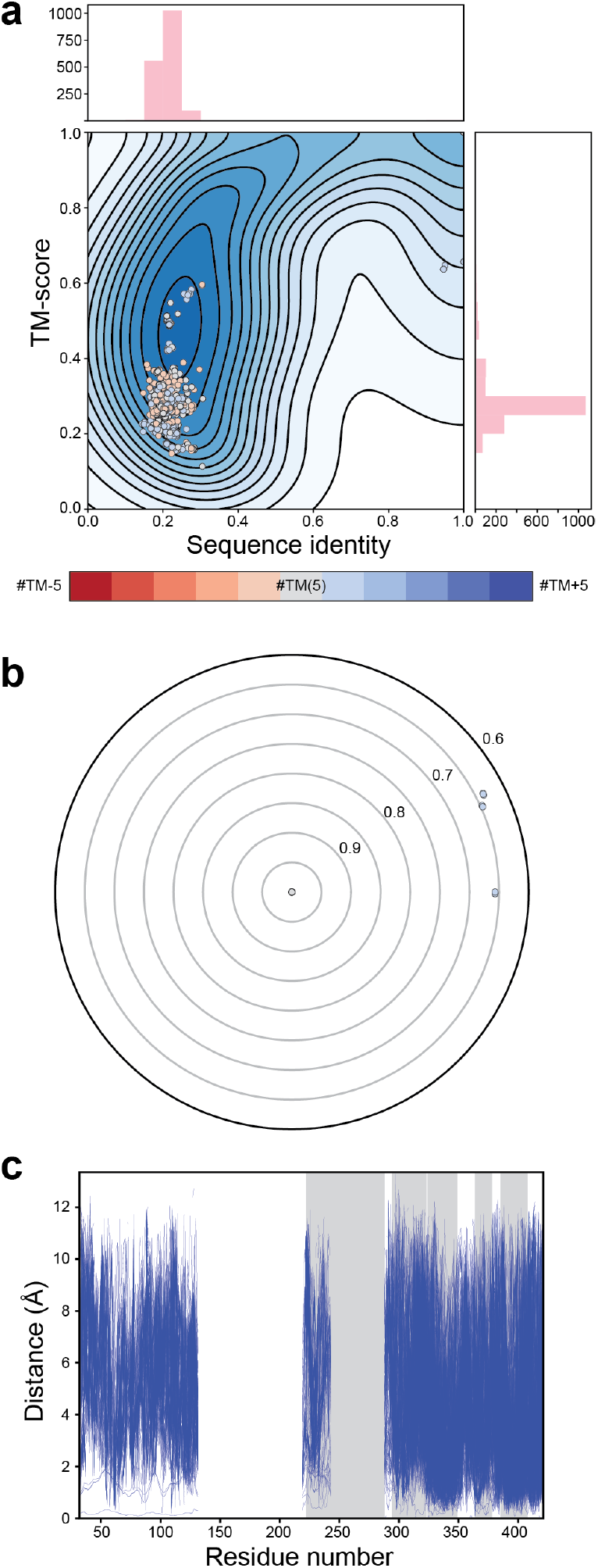
Plots provided to illustrate the network of sequence and structure relationships of a TM chain of interest, in this case one subunit of the Kv1.2 potassium channel (PDB code 2A79 chain J). a) Structure and sequence similarity for all pairwise alignments calculated for the structure of interest shown as points in a graph relating the sequence and structural similarity through the sequence identity and TM-score estimators. The histograms along the side and top report the distributions of the points in the graph, summed for each axis. Datapoints are colored based on the difference in the number of TM segments between the structure of reference and the structure being compared, following the legend, with bluer shades indicating that the structural neighbor has more TM segments, and redder shades indicating that the neighbor has fewer TM segments. b) Representation of the structural relationships between the structure of interest and its neighbors, as well as the similarity between those neighbors. All pairwise structure alignments calculated for the structure of interest with TM-score >0.6 are represented as points. The structure of interest is placed in the center of the plot. Distances between any two points in the graph are proportional to the TM-score distance between the two structures. Colors of individual points follow the legend. c) Regions of structural similarity and variability as a function of the protein sequence between the structural neighbors and the TM chain of interest. Ca-Ca distances are shown for each TM chain compared with the chain of interest. Grey regions indicate the TM regions in the structure of interest. Each blue line represents one comparison, and therefore, more dense regions indicate more structures with a given structural similarity in that region.

### 2.4 Symme try re cognition tools

By including complete descriptions of internal and quaternary symmetry in membrane-embedded segments of protein complexes, EncoMPASS provides a framework for assessing the effects of the membrane environment on the propensity of a protein to form symmetric structures. EncoMPASS includes the results of three different approaches for obtaining these descriptions. The first approach reports the output from two previously-developed symmetry detection tools, SymD (Kim *et al.*, 2010) and CE-Symm (Myers-Turnbull *et al.*, 2014; Bliven *et al.*, 2018), applied to the complexes as well as all individual TM chains. While a variety of methods have been designed for detecting symmetry or repeated elements in proteins (Mizuguchi and Gö, 1995; Murray *et al.*, 2004; Shih and Hwang, 2004; Abraham *et al.*, 2008; Guerler *et al.*, 2009), only SymD and CE-Symm overcome the challenge of extensive sequence divergence between internal repeats. Both methods achieve this by relying only on global structure-based alignments. In its most recent, unpublished version (1.61), SymD also reports a possible axis of symmetry. The user can filter out non-symmetric structures with the help of various cutoffs, default values of which were derived empirically by the authors. SymD does not report the range of residues that constitutes each of the repeats, whereas CE-Symm includes detailed information about the repeat ranges and their alignments. Furthermore, in its most recent version (Bliven *et al.*, 2018), CE-Symm can detect multiple symmetries simultaneously, provided that those symmetries relate sub-regions of the same symmetric repeats. In this way, CE-Symm can recognize dihedral or hierarchical symmetries (Fig. 5a). The program determines whether a candidate symmetry assignment should be “refined”, as well as its significance, based on structural similarity measured with the TM-score. The authors selected the corresponding thresholds so as to minimize false positives on a manually-curated set of 1,007 protein domains. A comparison between CE-Symm 1.0 and SymD 1.5b on this benchmark protein set demonstrated that the results from CE-Symm have a higher specificity compared to those obtained with SymD (Myers-Turnbull *et al,.* 2014).

**Figure 5.**
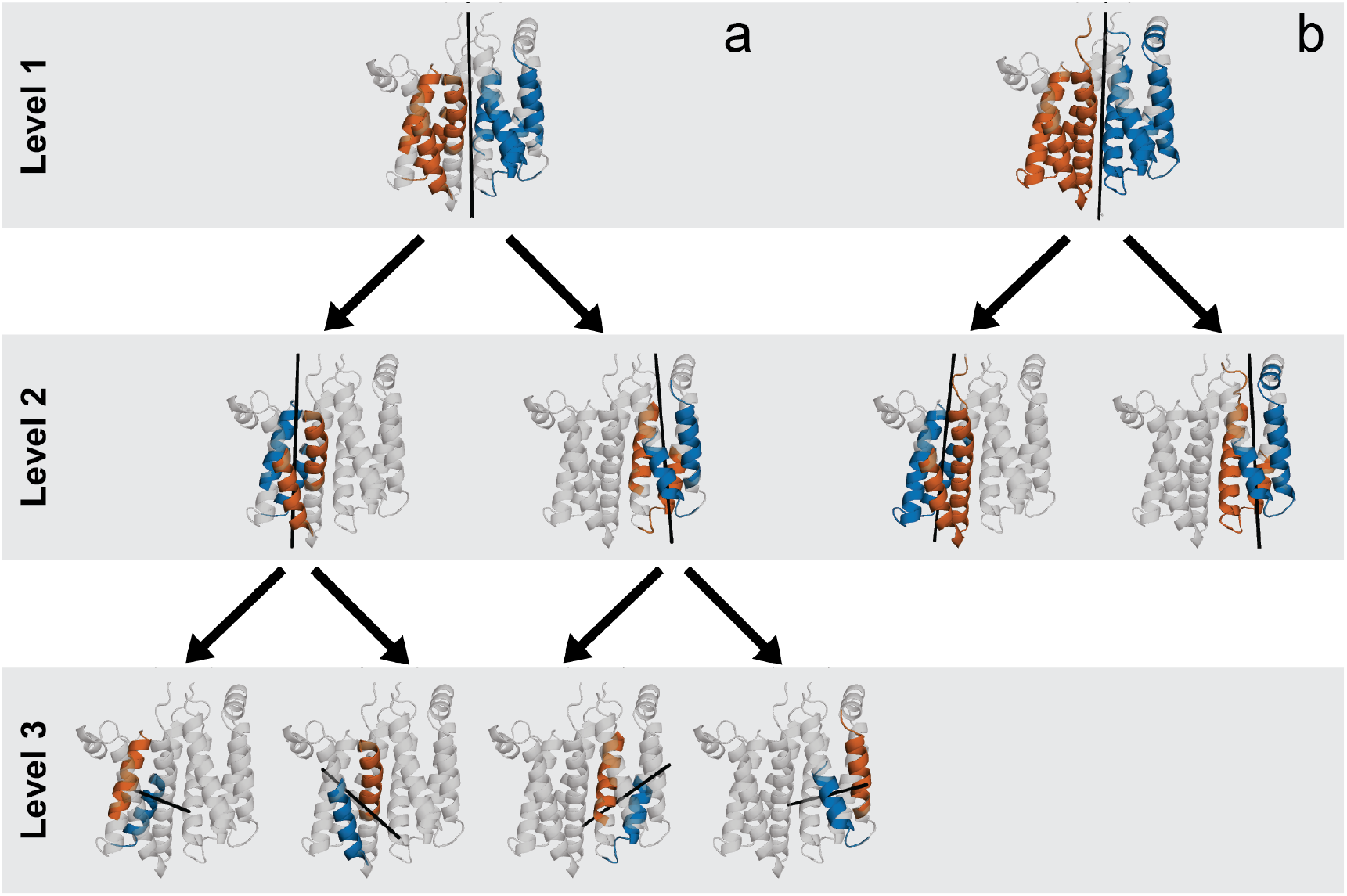
Multiple levels of symmetry detected in the PsrC subunit of polysulfide reductase (PDB code 2VPW, chain C). (a) CE-Symm recognizes three hierarchical levels of symmetry (highlighted by the gray background). Level 3 corresponds to the smallest building block present in all three levels of symmetry, and thereby also defines the maximum length of the repeats in levels 1 and 2. For PsrC, the building block is a single TM helix. However, the symmetry between two helices carries no functional significance. (b) Applying the CE-Symm-R procedure retains the symmetries described by levels 1 and 2, while eliminating those in level 3, which are unlikely to be functionally insightful due to their short length. The CE-Symm-R approach also results in a more extensive definition of the level 1 and 2 symmetric regions.

While CE-Symm assigns symmetries with high specificity, its developers estimated that due to its conservative thresholds, it underestimated the number of internally symmetric superfamilies in SCOP by about 27% (Myers-Turnbull *et al.*, 2014). However, the benchmark set on which CE-Symm was parametrized included only 30 membrane domains, or <3% of the whole set. Hence, we wanted to assess whether this estimate is equally valid for membrane proteins. We previously tested CE-Symm on a manually-curated benchmark set, MemSTATS, which includes complete symmetry descriptions for 97 a-helical membrane proteins and 22 β-barrel membrane proteins, each with distinct architecture (Aleksandrova *et al.*, 2018). Here we focus on the a-helical membrane proteins because they make up the majority of the membrane proteins and because the functional meaning of symmetry in the structure of β-barrel membrane proteins is poorly understood. For this a-helical subset, CE-Symm correctly identifies the presence and type of symmetry in about 80% of the TM chains and 74% of the complexes, although in some of these cases it finds only part of the symmetric repeat, i.e. the coverage is not complete (Fig. 6, Table 1). Notably, the rate of false-positive identifications of internal symmetry is about 14%, which is higher than the estimated rate of 3.3% on the benchmark set of Myers-Turnbull et al. In the following, we motivate and evaluate our strategy for overcoming the drawbacks of CE-Symm and for augmenting its capabilities with information obtained using SymD.

**Figure 6.**
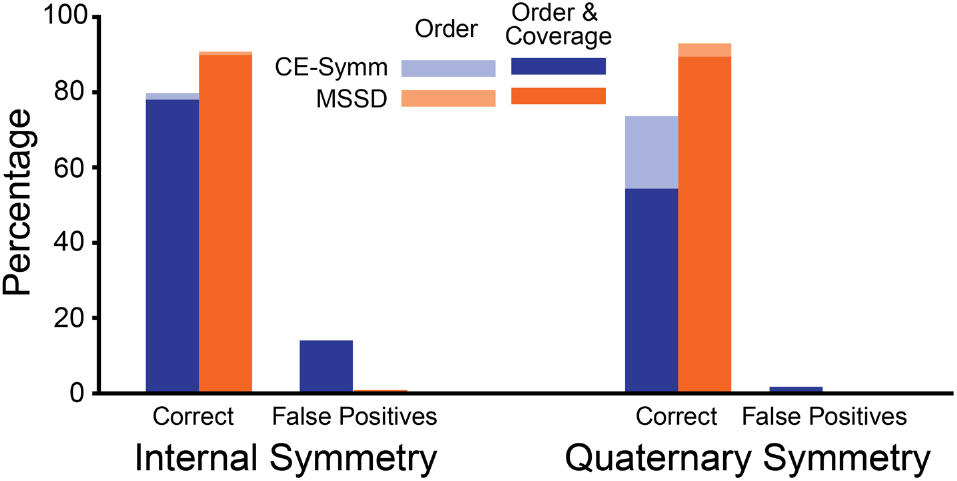
Symmetry detection results for CE-Symm (blue) and the multi-step symmetry detection method, MSSD used by EncoMPASS (orange) on a benchmark set of 97 alpha-helical proteins with distinct folds from MemSTATS. The methods are evaluated on the described symmetries within all TM chains in the set (internal symmetry) and within all complexes with >1 TM chain (quaternary symmetry): in total, 114 and 57 symmetry descriptions, respectively. The percentage of symmetries identified correctly, which is the sum of the true-positive and true-negative rates (Table 1), as well as the false-positive rate are shown for each method. In cases where a method detects the symmetry described in the benchmark, but reports a symmetry repeat that is only part of the benchmark repeat (i.e. it is missing at least 20 consecutive residues), the correct rate is shown in lighter color.

**Table 1:** Symmetry assignment benchmarking results

**a) Correct symmetry assignments including True Positives and True Negatives**
| Subset | Total<sup>a</sup> | True Positives |  |  | True Positives<br>with Complete Coverage<sup>b</sup> |  |  | True Negatives |  |  |
| --- | --- | --- | --- | --- | --- | --- | --- | --- | --- | --- |
|  |  | CE-Symm | MSSD | MSSD+I | CE-Symm | MSSD | MSSD+I | CE-Symm | MSSD | MSSD+I |
| Chains | 114 | 29 | 37 | 41 | 27 | 36 | 40 | 62 | 67 | 67 |
| Complexes | 57 | 39 | 50 | - | 28 | 48 | - | 3 | 3 | - |

| Subset | Total | False Positives |  |  | False Negatives |  |  |
| --- | --- | --- | --- | --- | --- | --- | --- |
|  |  | CE-Symm | MSSD | MSSD+I | CE-Symm | MSSD | MSSD+I |
| Chains | 114 | 16 | 1 | 1 | 23 | 10 | 6 |
| Complexes | 57 | 1 | 0 | - | 15 | 4 | - |

Results are reported as number of symmetry descriptions for each subset in the MemSTATS dataset, compared to the total number of descriptions<sup>a</sup>. Results were obtained using CE-Symm or using the MSSD method in EncoMPASS either without inferring symmetry from related structures (MSSD), or after inferring symmetry from related structures (MSSD+I).<sup>b</sup>Symmetries assignments in which the extent of the coverage is correctly assessed, in addition to the symmetry order. Chains: results for all α-helical membrane-spanning chains, i.e. with internal symmetry. Complexes: α-helical complexes with >1 membrane-spanning chain, i.e. quaternary symmetry.

### 2.5. Symmetry detection in membrane protein chains

Given the potential mentioned above that CE-Symm could fail to detect difficult symmetries in a set of membrane protein structures, an obvious strategy would be to replace the default thresholds with more lenient values. However, this approach could also increase the number of false positives. Fortunately, the lipid bilayer creates natural constraints that can be used to help filter out such false positives. Specifically, in α-helical TM proteins, the symmetry between a single TM helix and another TM helix can be considered a trivial observation with no functional significance. Therefore, by ignoring symmetries between repeats that each comprise less than two TM segments, we can eliminate such cases. As mentioned, this filter allows us to screen more permissive approaches for detecting possible symmetries and to consequently find more true positives with a potentially more comprehensive and functionally relevant definition of the symmetric repeats. Note that this approach has so far been tailored to α-helical TM proteins, which constitute the majority of membrane proteins, but could also readily be extended to TM β-barrels in the future.

The first screen that we applied was to process each TM chain with CE-Symm customized by halving the TM-score threshold for refinement while also doubling the window size of aligned fragment pairs considered during the structural alignment. We refer to these modifications collectively as CE-Symm-L, for “lenient”.

The CE-Symm-L strategy detects divergent global symmetries but provides no mechanism for identifying two distinct symmetries within a single structure that cannot be related to one another, such as that found in GltPh (Fig. 7). We therefore flag any structures in which CE-Symm-L has detected symmetry, but in doing so omitted one or more large continuous fragments of the protein. Such fragments are extracted and re-processed with CE-Symm-L. A similar strategy is also applied to results from SymD. Specifically, we examine the best self-alignment produced by SymD for large fractions of a protein chain that SymD has left unaligned or for which no symmetry was reported by CE-Symm. We then extract these fragments and process them separately with CE-Symm to assess the plausibility of symmetry existing therein, as well as to obtain more detailed information about the repeats.

**Figure 7.**
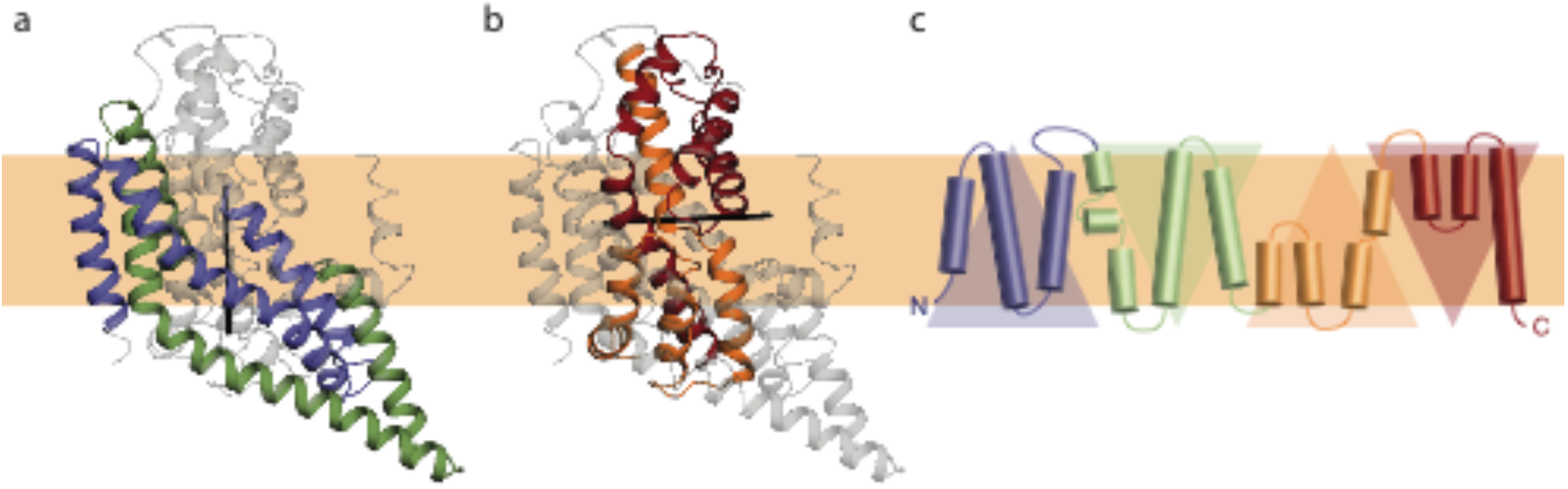
The bacterial glutamate transporter homolog GltPh (PDB code 2NWX chain A) contains two small, non-hierarchical symmetries within each protomer, which cannot be captured by CE-Symm using default parameters. The first half of the protomer (a) is C2-pseudo-symmetric, as is the second half of the protomer (b), but the fold of the first two repeats is too distinct from the fold of the other two repeats (c) for CE-Symm to identify any connection. Because the CE-Symm algorithm, by construction, cannot report the two symmetry relationships simultaneously, and because each of the individual symmetries alone scores below the default TM-score threshold, CE-Symm detects no symmetry in this structure.

The next processing step addresses an issue inherent to the hierarchical strategy used by CE-Symm. Specifically, this strategy occasionally finds additional axes and levels of symmetry by breaking down functionally meaningful repeats into elements that are too small to be relevant (Fig. 5a). The aforementioned filtering procedure will eliminate these results entirely, thus discarding potentially important information about the relationship between the larger repeats. To correct for this, we identify all such possible results produced by CE-Symm or CE-Symm-L. We then reprocess these structures with CE-Symm, but this time limiting the maximum number of symmetry levels allowed. This in effect forces CE-Symm to report the relationships detected between the larger fragments of the protein. We call this procedure CE-Symm-R, for “restrictive” (Fig. 8b).

**Figure 8.**
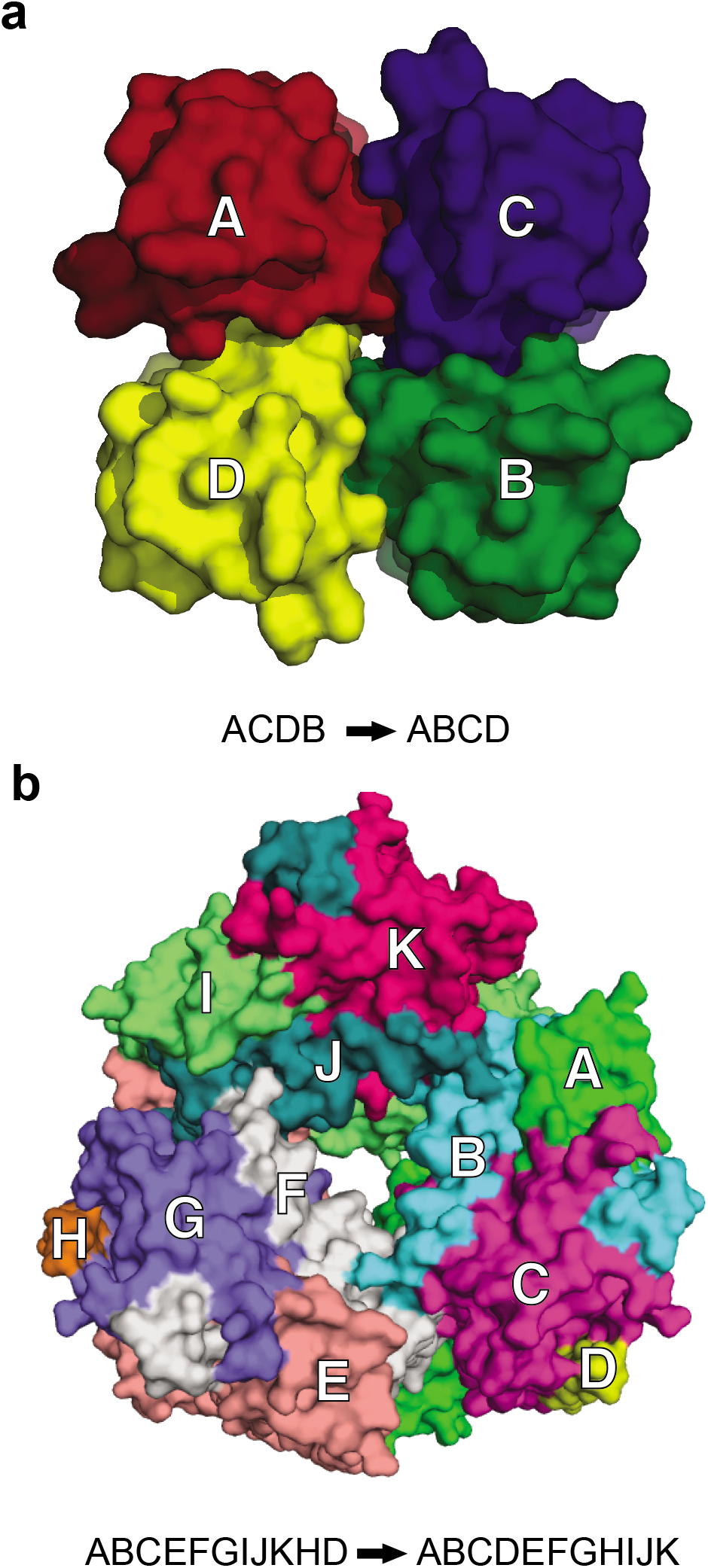
Strategies for ordering the chains of a complex to aid detection of their symmetric arrangements. (a) The PDB coordinates of the KcsA channel (PDB code 1R3J) present the chains in an order that is not conducive to detecting their symmetric arrangement because there is no single transformation that can match all subunits. To address this issue, the chains can be re-arranged starting from a chain at one end of the complex and progressively adding the closest unordered neighbor. Repeating this until all chains have been accounted for allows for the C4 quaternary symmetry to be detected. (b) Particulate methane monooxygenase (PDB code 3RFA) contains a C3 symmetry relating chains A-C, E-G and I-K, which the previous strategy fails to detect because chains D and H have no symmetry equivalent. Instead, a SymD self-alignment can be used for re-arranging the order of the chains so that the symmetry can be detected.

Two additional procedures were implemented that restrict specific parameters during CE-Symm processing, called CE-Symm-RMSD and CE-Symm-O. In the CE-Symm-RMSD procedure, we restrict the maximum RMSD allowed between repeats during the refinement step of the CE-Symm algorithm, which is particularly useful when the protein is composed of many parallel alpha-helices.

In the second procedure, CE-Symm-O (for “order”), we address cases in which SymD identifies a higher symmetry order than CE-Symm (Myers-Turnbull et al., 2014), either because CE-Symm has not identified enough repeats or because CE-Symm has used multiple hierarchical levels to describe the same symmetry relationships. To this end, we scan the results from CE-Symm and SymD and flag cases in which the number of repeats identified by SymD is a multiple of the number identified by CE-Symm, and where the symmetry detected by SymD has order >3. These flagged structures are reprocessed with CE-Symm, but with a parameter that enforces the search only to results corresponding to the order detected by SymD.

By combining these four procedures (CE-Symm-L, -R, -RMSD and -O), we obtain multiple possible assignments for the symmetries in each protein chain. After filtering out trivial assignments with <2 TM crossings in a repeat and <4 TM crossings altogether, we consolidate the results into a single symmetry assignment by imposing the following rules: the optimal symmetry is that with the greatest number of symmetry levels, thus favoring, e.g., dihedral symmetries over cyclical ones; if all strategies report the same symmetry level, however, we select the result with the largest number of symmetrically aligned residues overall (see STAR Methods).

Assessing our approach on the a-helical membrane proteins in the MemSTATS symmetry benchmark set (Fig. 6, Table 1), we note that the rate of true positives among internal symmetries increases by >11%, while the rate of false positives has decreased sixteenfold.

### 2.6. Symmetry detection in membrane protein complexes

While CE-Symm and SymD were designed with internal symmetries in mind, their algorithms can readily be applied to detecting quaternary symmetry as well. The main obstacle for this endeavor is that the algorithms fail to detect even perfect symmetry if the chains in the protein coordinate file are not ordered appropriately (Fig. 8). Therefore, if a complex has more than three membrane-spanning chains, we apply three strategies to guess a more suitable order for their chain names. First, we order the chains starting from a peripheral chain and progressively add the closest unaccounted neighboring chain. This strategy works well in most cases, but it is ineffective for more complicated complexes, such as the methane monooxygenase structure 3RFA (Fig. 8b). Thus, as a second strategy, we use the matching of chains found in the best self-alignment produced by SymD to guess the optimal order. This approach also allows a subset of chains to be considered thereby allowing symmetric relationships in a comparatively small portion of a large complex to be detected. Finally, we combine the two strategies consecutively to provide a third guess of the most suitable ordering of chains.

The complex with each of the four possible chain orders (including the original one from the Protein Databank) is then processed with CE-Symm, SymD, CE-Symm-L, CE-Symm-R, CE-Symm-RMSD, and CE-Symm-O. Similar filtering and selection procedures to those used for internal symmetry are applied to identify all possible symmetries at the quaternary level. Overall, our approach identifies correctly 93% of the quaternary symmetries of the a-helical membrane protein cases in the MemSTATS benchmark set, compared to the 74% identified correctly by CE-Symm alone (Fig. 6).

### 2.7. Inference of symmetric repeats from related structures

Our approach thus far has aimed at optimizing the detection of quaternary and internal symmetry in a single membrane protein structure. However, many membrane proteins undergo large conformational changes to accomplish their function. During these changes, the relationship between internal repeats might change noticeably, making them more or less symmetric, and these changes in symmetry can also relate to the functional mechanisms of the protein (Forrest, 2015). Therefore, it is useful to track such changes between available structures of the same protein in different states. Furthermore, we want to be able to compare how repeats differ between proteins that are structurally similar (TM-score ≥0.6 as defined in Section 2.4). To this end, we developed a strategy to conserve the definition of repeats across related structures, taking advantage of the similarities between chains identified in the structural alignment analysis.

If symmetry has been detected in any of the chains defined as similar (either structurally or by sequence) to the chain of interest, then the symmetric repeats of the former are mapped on the residues of the latter with the help of the structure-based alignment. In cases where there are multiple related chains in which symmetry has been detected, the assignment with the highest symmetry level and/or the highest coverage is used as the reference. To quantify the structural relationships between the newly-identified repeats, we implement the same procedure as the one used by CE-Symm.

Using these strategy for inference of symmetric repeats together with our multi-step symmetry detection (MSSD) approach results in 95% correct identifications of the internal symmetries of the a-helical membrane proteins in the MemSTATS benchmark set, without changing the low rate of false-positives (Table 1).

## Discussion

In order to meet the need for a clearer overview of the structural relationships and symmetry properties of membrane proteins, the EncoMPASS database combines structural information about membrane proteins taken from the OPM, PDB, and PDBTM databases, and makes them available through a graphical interface. Due to the requirement for high-quality data, only X-ray structures with resolution ≤3.5 Å are considered, for a total of 2344 entries, which constitutes ~67% of the polytopic structures in OPM. The three databases do not always concur on all the characteristics of the membrane proteins they describe. Notably, the proposed biological assembly often differs. In these cases we applied a methodology for selecting the most reliable description of the assembly. Of the initial 2400 structures, 1345 were thus corrected. Although it was not possible to quantify precisely the number of residual errors in the definition of biological units, we expect that such errors have been reduced substantially by comparative analyses and structural consistency checks.

Once a set of revised and reliable structures had been compiled, the first task of EncoMPASS was to identify structural relationships between them. The brute-force strategy of comparing all possible pairs would be very time-consuming and likely to produce a large number of false positives. Our strategy of comparing structures according to simple fixed rules based on transmembrane topology and soluble domain size eliminates nearly 84% of all possible comparisons, and therefore greatly reduces the number of required calculations. More importantly, these rules reduce the number of false-positive structural relationships, while not being overly restrictive regarding the topological similarity of the two chains. As with any set of rigid rules, this strategy is a compromise that could result in some false negatives. For example, structures of the SWEET and SemiSWEET transporters are not included in the set of allowed comparisons. Thus, despite their convenience, these rigid rules may be unable to fully reflect the complexity of the network of structural relationships: in the future, we plan to replace this procedure with a more versatile approach.

Another concern regarding the detection of structural relationships is the robustness of the structural alignment methodology, which motivated the choice to limit the structural resolution to 3.5 Å and to consider only structures determined by X-ray crystallography. As more structures become available from methods such as cryo-electron microscopy and as the reliability of the modeling of such low-resolution structures improves, we anticipate that it will be possible to loosen this constraint.

The addition of new structures in the future will potentially present new challenges for the symmetry detection and analysis methods, which currently consider a set of empirically-derived parameter combinations that may not be sufficient for unforeseen reasons. Therefore, as the database expands, it may be necessary to implement additional processing steps to improve our assessment of the symmetries. In addition, newly developed symmetry detection algorithms, such as the method used to curate quaternary symmetries in the PDB database, can be evaluated and incorporated within the framework of the MSSD analysis.

As mentioned above, symmetry and pseudo-symmetry can have important implications for understanding the folding, function, and evolution of membrane proteins. For example, well-defined asymmetry can be central to the function of many secondary active transporters (Forrest 2015). The detection of such asymmetries, which may reflect concerted rearrangements of groups of helices, in some cases may require the development of a new set of definitions and procedures. This will be the focus of future work.

In conclusion, EncoMPASS offers a systematic, automated, and freely-available library of membrane protein structures and relationships based on a consistent set of similarity estimators and symmetry descriptions. We therefore expect that EncoMPASS will prove useful not only for the user interested in specific proteins, but also for more general studies of the nature of membrane protein structure and evolution.

## Acknowledgements

We thank the CE-Symm and SymD developers for providing unpublished versions of the code and helpful discussions, Wim Vranken, Gabriele Orlando and Daniele Raimondo for interesting conversations on structural and topological alignment techniques, Philippe Youkharibache for pointing out the usefulness of the maxrmsd parameter in CE-Symm, Amarendra Yavatkar and Srujan Ganta (NINDS) for building the web site, the NHBLI LOBOS administrators for technical assistance, and Marcus Stamm for useful discussions.

## STAR Methods

### 2.8 WORKFLOW

#### Initial structure collection

We chose the OPM database (Lomize *et al.*, 2012) as the primary source of information on which EncoMPASS is built for the following reasons: its entries are manually curated, which generally improves accuracy, the insertion of the membrane proteins in the model lipid bilayer is accurate (Shimizu *et al.*, 2018), and it contains information on the transmembrane segments of each TM chain. Nevertheless, manual curation can, at times, produce errors, and thus, the corresponding structure is also downloaded from the PDB (Berman *et al.*, 2000). The EncoMPASS procedure parses both files and chooses which to rely on, depending on whether they contain errors or omissions in the PDB format, such as repeated or disordered residue indices in a chain, or any anomaly resulting in a parsing error. In cases where both files are acceptable, preference is given to the OPM structure. Even after this step, the PDB and OPM files are both preserved, for reference in case errors are found in subsequent steps.

#### Uniform structure parsing

Coordinate files from OPM and PDB are converted to a uniform EncoMPASS PDB format, for compatibility with the structure analysis software:

- Non-standard residues and “UNK” entries are deleted. The only exception is MSE residues, which are converted to MET;
- ANISOU, MASTER, and CONECT lines are deleted;
- Chains are parsed with BioPython, which is employed in later steps of the procedure. If BioPython returns an error, then the structure will be scheduled for correction (see “Biological assembly correction strategies” section);
- DUM entries, dummy atoms used by OPM to delineate the membrane boundaries, are preserved;
- Residues with incomplete backbones are deleted;
- For entries that assign alternate conformations or locations (AltLocs) for any of the residues, the average occupancy is calculated over all atoms assigned to a given alternate conformation (ATOM entries with the same character in column 17). The conformation with the maximum average occupancy is chosen, atoms in the other conformations are deleted, and the AltLoc character is reset to empty.

#### Strategy for correcting biological assemblies

We found a number of PDB entries in which the instructions for generating the biological unit were missing or misleading. Misleading instructions can be especially problematic for predicting the correct insertion of the complex in the bilayer. Many techniques to solve this class of problems exist. We follow the strategy used in PDBTM (Kozma *et al.*, 2013), in which structures are first divided into a “safe” set that can be used as a reference, and a set of potentially problematic structures. A structure is considered potentially problematic if:

- One or more chains described in the ATOM lines have identical sequences;
- There is only one chain, but there is no indication of any geometrical transformation to obtain a biologically active complex;
- The number of subunits multiplied by the number of their geometrical transformations stated in the instructions of the PDB file does not match the number of subunits found in the OPM file;
- At least one of the geometrical transformations results in a translation whose axis is parallel to the axis of rotation;
- The geometrical transformations produce an assembly in which chains are detached.

For each structure in the potentially problematic set, we search the “safe” set for the structure with the highest sequence similarity, provided it is >90%. If such a structure is found, we analyze its biological assembly and apply the same transformations to the problematic structure. In case no similar structure is found, we download the corresponding coordinate file from the PDBTM database. However, the PDBTM database only reports the transmembrane portion of the complexes in their correct biological assembly: to recover the full complex we therefore apply a set of partial structure superpositions.

#### Biological assembly correction test cases

Thirteen structures were chosen as test cases for the effectiveness of the biological assembly correction strategy. Structures with incorrect BIOMATRIX transformations are readily detected by visual inspection, whereas erroneous oligomerization states can be difficult to identify, as they depend on the function of the protein complex. Structures with incorrect BIOMATRIX transformations: 1QJ9, 2A79, 2B6O, 2VQI, 3DH4, 3LNM, 3M9I, 3QQ2, 3RB2, 4JRE, 4R50. Structures with incorrect oligomerization states: 1F88, 2ATK.

#### Insertion in the model lipid bilayer

Once all checks have been performed and the coordinate files reflect the complete biological assembly, the structures that were modified or taken from either PDB or PDBTM are inserted in the model lipid bilayer with PPM (Lomize *et al.*, 2012), the algorithm used by OPM. Once inserted, each TM chain of each structure is analyzed to compute the number and extent of the TM regions. We define a TM region as a polypeptide segment comprising >2 consecutive residues from a TM segment as defined by PPM. Since PPM TM segments are defined as secondary structure elements that are at least partially inside the membrane, this rule ensures that all defined TM regions contain a minimum secondary structure content, while avoiding short or shallow membrane regions. By allowing more than one PPM TM segment to belong to a single TM region, we also avoid double counting of reentrant regions (present in 46% of the structures of our database) and of TM segments with unwound regions separating two or more secondary structure elements.

PPM has a tendency to place the boundaries of the model lipid bilayer so that small loops connecting membrane-spanning helices end up inside the membrane. The aforementioned definition of a TM region would categorize two such helices as only one TM region, even though such an assignment arguably would not reflect the correct topology. To avoid this issue, therefore, we define the membrane as 2 Å thinner than that considered by PPM, i.e. with boundaries parallel to the layers defined by PPM but with each boundary displaced by 1 Å towards the interior of the membrane.

#### Selection rules for performing a structure alignment

To avoid comparisons between structures that are obviously topologically unrelated, we apply the following selection rules to determine whether two TM chains should be aligned.

- For single-spanning TM chains, both chains must contain either no significant terminal domains (>100 amino acids in length), or at least one extramembrane domain of similar length, defined as at least half the size of the longest terminal segment.
- For chains with 2 TM regions, the same rule applies as for 1-TM segments. In addition, we require that any domains between the two TM regions be of similar length.

These rules become impractical and ineffective as the number of TM regions increases. Thus, for chains with 3 or more TM segments, we introduce a new set of rules:

- Two chains containing the same number of TM regions are compared;
- Two chains with different numbers of TM regions are only be compared if the chain with the smaller number of segments has at least 75% as many TM segments as the other chain.

Comparisons between a-helical and β-stranded TM chains are never allowed, whereas comparisons between two β-stranded TM chains are always allowed.

#### Structure and sequence alignments

Structure alignments are performed with Fr-TM-Align, which returns the RMSD and TM-score values calculated on the aligned Cα atoms, and the corresponding sequence alignment. We also calculate a structure-independent sequence alignment with MUSCLE. In this case, instead of considering the sequence of amino acids in the structure file, which only includes those residues for which density could be resolved, we take the sequence deposited in the PDB, which is the construct used for crystallization. For both these sequence alignments we calculate the sequence identity normalized to the number of matches/mismatches (i.e., without taking gaps into account).

#### Symmetry detection with default parameters of CE-Symm and SymD

To analyze the symmetry in each structure, we process each TM chain and complex in EncoMPASS with SymD version 1.61 and CE-Symm version 2.0-RC2 using default parameters. CE-Symm relies on a random seed, which in some cases affects its ability to detect symmetry. Therefore, for each procedure involving CE-Symm, the program is run with three different seeds, namely 3, 5 and 10. In cases where this repetition creates variations in the results, we choose the outcome containing a symmetry with repeat length of >40 amino acids, the highest number of repeats, and the largest number of amino acids.

#### Multi-step approach for symmetry detection

Multiple screens and customized parameters are applied in parallel to extract additional symmetry assignments from CE-Symm and SymD. We refer to the following procedures:

- CE-Symm-L: CE-Symm with options –unrefinedscorethreshold = 0.2, –winsize = 16;
- CE-Symm-R: adds the option –symmlevels = (*k* – 1), where *k* is the number of symmetry levels in a result from default CE-Symm or CE-Symm-L, in which the length of the repeats is <2 TM crossings;
- CE-Symm-O: adds the option –order = k, where *k* = |360°/θ| and θ is the unit angle describing the symmetry reported by SymD;
- CE-Symm-RMSD: CE-Symm-L with the additional option –maxrmsd = k, where *k* is 2.5 or 3.

We also break TM chains into fragments, when appropriate, where a fragment must be continuous and contain >80 amino acids, corresponding to roughly four TM helices. Each fragment is then processed with CE-Symm and SymD with default options.

For each TM chain-ordered configuration of a complex (see Results) and for each of its TM chains, the symmetry is analyzed by running: (i) SymD; (ii) CE-Symm with default options; (iii) CE-Symm-L; (iv) CE-Symm-R; (v) CE-Symm-O; and (vi) CE-Symm-RMSD, as appropriate. We then compare the results from each procedure and select an optimal result for each chain and complex, as described in the Results section.

#### Inference of symmetric repeats

For inferring symmetry from one chain to another, we first rank the results from the MSSD approach for all TM chains, from best to worst according to the following rules: 1) higher order is preferred, e.g., D10 > D2 > C2 > R, where R stands for repeated or open symmetry; 2) if the order is the same, the result with higher number of aligned residues is preferred; 3) if both the order and the aligned length are the same, the result obtained with the structure with the most residues is preferred. Using these criteria, for each TM chain, we select the highest-ranking result of all TM chains that are either structurally-related (TM-score > 0.6) or sequence-related (identity > 0.85) and use it as a template. We use the sequence alignment between the two chains to infer the corresponding residues in the target chain. All identified repeats in the target structure are superimposed onto the first repeat in that structure, using symmetry operators calculated for each symmetry level with a Kabsch algorithm (Kabsch, 1976; Kabsch, 1978). For each pair of repeats, the RMSD and TM-score representing their structural relationship are calculated and, where appropriate, the average over all pairs is reported.

### 2.9 CODE AVAILABILITY

EncoMPASS is created and updated automatically through a library of Python 3.5 routines, freely available at https://github.com/EncoMPASS-code/EncoMPASS. The bundle does not contain the external programs for finding the correct orientation of the protein in the membrane (PPM (Lomize *et al.*, 2012)), for sequence alignment (MUSCLE (Edgar, 2004)), for structure alignment (Fr-TM-Align (Pandit and Skolnick, 2008)), or for symmetry detection (CE-Symm (Myers-Turnbull *et al.*, 2014) and SymD (Kim *et al.*, 2010)), but these can all be obtained freely.

### 2.10 WEBSERVER

The EncoMPASS database web server is hosted at https://encompass.ninds.nih.gov. From the home page the user can navigate to any entry by entering the corresponding PDB code in the search bar. The online database is composed of a set of webpages, each describing a single membrane protein complex. All individual TM chains are also described in separate webpages that can be accessed either from the webpage of the related complex or directly from the search bar, by specifying the name of the chain (e.g. ‘4hea_C’), and contain information unique to individual chains, such as structural relationships with other TM chains in the database and internal symmetries. The server is described in detailed elsewhere (Sarti *et al.*, in preparation).

### 2.11 DOWNLOADABLE MATERIAL

The entire EncoMPASS database can be downloaded at https://encompass.ninds.nih.gov/downloads. In the bundle, all displayed coordinate files are included, as well as all the sequence and structure similarity estimators used to produce the graphs, i.e., TM-score, RMSD, structure-wise SeqID, and sequence-wise SeqID.

## Abbreviations

TM: transmembrane

## References

Abraham, A.L. et al. (2008) Swelfe: A detector of internal repeats in sequences and structures. Bioinformatics, 24, 1536–1537.

Aleksandrova, A.A., Sarti, E., and Forrest, L.R. (2018). MemSTATS benchmark v1.0. doi: 10.5281/zenodo.1345122.

Andreeva, A. et al. (2007) Data growth and its impact on the SCOP database: new developments. Nucleic Acids Res., 36, D419–D425.

Balaji, S. (2015) Internal symmetry in protein structures: prevalence, functional relevance and evolution. Curr. Opin. Struct. Biol., 32, 156–66.

Berezovsky, I.N. et al. (2016) Basic units of protein structure, folding, and function. Prog. Biophys. Mol. Biol.

Berman, H.M. et al. (2000) The Protein Data Bank. Nucleic Acids Res., 28, 235–242.

Bliven, S.E. et al. (2018). Analyzing the symmetrical arrangement of structural repeats in proteins with CE-Symm. bioRxiv, 1–18.

Bork, P. (1991) Shuffled domains in extracellular proteins. FEBS Lett., 286, 47–54.

Bourne, P.E. and Shindyalov, I.N. (2003) Structure comparison and alignment. In, Bourne, P.E. and Weissig, H. (eds), Structural Bioinformatics, Volume 44. John Wiley & Sons, Inc., pp. 321–337.

Bowie, J. (2001) Stabilizing membrane proteins. Curr. Opin. Struct. Biol., 11, 397–402.

Bull, S.C. and Doig, A.J. (2015) Properties of protein drug target classes. PLoS One, 10, e0117955.

Chen, H. et al. (2009) A simple method of identifying symmetric substructures of proteins. Comput. Biol. Chem., 33, 100–107.

Choi, S. et al. (2008) Common occurrence of internal repeat symmetry in membrane proteins. Proteins, 71, 68–80.

Cournia, Z. et al. (2015) Membrane protein structure, function, and dynamics: a perspective from experiments and theory. J. Membr. Biol., 248, 611–640.

Davey, J. (2004) G-protein-coupled receptors: new approaches to maximise the impact of GPCRs in drug discovery. Expert Opin. Ther. Targets, 8, 165–170.

Dib, L. and Carbone, A. (2012) Protein fragments: functional and structural roles of their coevolution networks. PLoS One, 7, e48124.

Duran, A.M. and Meiler, J. (2013) Inverted topologies in a membrane: a mini-review. Comput. Struct. Biotechnol. J., 8, 1–8.

Edgar, R.C. MUSCLE: multiple sequence alignment with improved accuracy and speed. In, Proceedings. 2004 IEEE Computational Systems Bioinformatics Conference, 2004. CSB 2004. IEEE, pp. 689–690.

Espada, R. et al. (2015) Repeat proteins challenge the concept of structural domains. Biochem. Soc. Trans., 43, 844–849.

Forrest, L.R. (2015) Structural symmetry in membrane proteins. Annu. Rev. Biophys., 44, 311–337.

Frishman, D. (2010) Structural Bioinformatics of Membrane Proteins Springer Vienna, Vienna.

Golmohammadi, S.K. et al. (2007) Classification of cell membrane proteins. In, 2007 Frontiers in the Convergence of Bioscience and Information Technologies. IEEE, pp. 153–158.

Goodsell, D.S. and Olson, A.J. (2000) Structural symmetry and protein function. Annu. Rev. Biophys. Biomol. Struct., 29, 105–53.

Guda, C. et al. (2001) A new algorithm for the alignment of multiple protein structures using Monte Caro optimization. Pacific Symp. Biocomput., 6, 275–286.

Guerler, A. et al. (2009) Symmetric structures in the universe of protein folds. J. Chem. Inf. Model., 49, 2147–2151.

Holm, L. and Sander, C. (1995) Dali: a network tool for protein structure comparison. Trends Biochem. Sci., 20, 478–480.

Hubbard, T.J.P. et al. (1997) SCOP: a Structural Classification of Proteins database. Nucleic Acids Res., 25, 236–239.

Janin, J. and Wodak, S.J. (1983) Structural domains in proteins and their role in the dynamics of protein function. Prog. Biophys. Mol. Biol., 42, 21–78.

Kabsch, W. (1976). A solution for the best rotation to relate two sets of vectors. Acta Cryst. Sect. A 32, 922–923.

Kabsch, W. (1978). A discussion of the solution for the best rotation to relate two sets of vectors. Acta Cryst. Sect. A 34, 827–828.

Kim, C. et al. (2010) Detecting internally symmetric protein structures. BMC Bioinformatics, 11, 303.

Levy, E.D. et al. (2006) 3D complex: a structural classification of protein complexes. PLoS Comput. Biol., 2, 1395–1406.

Liu, Y. et al. (2004) TM protein domains rarely use covalent domain recombination as an evolutionary mechanism. Proc. Natl. Acad. Sci., 101, 3495–3497.

Lomize, A.L. et al. (2006) Positioning of proteins in membranes: a computational approach. Protein Sci., 15, 1318–1333.

Lomize, M.A. et al. (2006) OPM: orientations of proteins in membranes database. Bioinformatics, 22, 623–625.

Lomize, M.A. et al. (2012). OPM database and PPM web server: resources for positioning of proteins in membranes. Nucleic Acids Res. 40, D370–D376.

Mizuguchi, K. and Gö, N. (1995) Comparison of spatial arrangements of secondary structural elements in proteins. Protein Eng. Des. Sel., 8, 353.

Moraes, I. et al. (2014) Membrane protein structure determination — the next generation. Biochim. Biophys. Acta - Biomembr., 1838, 78–87.

Murray, K.B. et al. (2004) Toward the detection and validation of repeats in protein structure. Proteins Struct. Funct. Genet., 57, 365–380.

Myers-Turnbull, D. et al. (2014) Systematic detection of internal symmetry in proteins using CE-symm. J. Mol. Biol., 426, 2255–2268.

Neumann, S. et al. (2010) Current status on membrane protein structure classification. Proteins Struct. Funct. Bioinf., 78, 1760–1773.

Orengo, C. et al. (1997) CATH – a hierarchic classification of protein domain structures. Structure, 5, 1093–1109.

Pandit, S. and Skolnick, J. (2008) Fr-TM-align: a new protein structural alignment method based on fragment alignments and the TM-score. BMC Bioinformatics, 9, 531.

Petrey, D. and Honig, B. (2009) Is protein classification necessary? Toward alternative approaches to function annotation. Curr. Opin. Struct. Biol., 19, 363–368.

Prilusky, J. (1996) OCA, a browser-database for protein structure/function.

Richardson, J.S. (1981) The anatomy and taxonomy of protein structure. In, Advances in Protein Chemistry. Elsevier Inc., pp. 167–339.

Saier, M.H. (2006) TCDB: the Transporter Classification Database for membrane transport protein analyses and information. Nucleic Acids Res., 34, D181–D186.

Shih, E.S.C. et al. (2006) OPAAS: A web server for optimal, permuted, and other alternative alignments of protein structures. Nucleic Acids Res., 34, 95–98.

Shih, E.S.C. and Hwang, M.J. (2004) Alternative alignments from comparison of protein structures. Proteins Struct. Funct. Genet., 56, 519–527.

Shimizu, K. et al. (2018) Comparative analysis of membrane protein structure databases. BBA Biomembrane, 1860(5), 1077–1091.

Shindyalov, I.N. and Bourne, P.E. (1998) Protein structure alignment by incremental combinatorial extension (CE) of the optimal path. Protein Eng. Des. Sel., 11, 739–747.

Skolnick, J. et al. (2009) The continuity of protein structure space is an intrinsic property of proteins. Proc. Natl. Acad. Sci., 106, 15690–15695.

Sojo, V. et al. (2016) Membrane proteins are dramatically less conserved than water-soluble proteins across the tree of life. Mol. Biol. Evol., 33, 2874–2884.

Stamm, M. and Forrest, L.R. (2015) Structure alignment of membrane proteins: accuracy of available tools and a consensus strategy. Proteins Struct. Funct. Bioinforma., 1720–32.

Stansfeld, P.J. et al. (2015) MemProtMD: automated insertion of membrane protein structures into explicit lipid membranes. Structure, 23, 1–12.

Stansfeld, P.J. and Sansom, M.S.P. (2014) MemProtMD: membrane protein structures and simulations. Biophys. J., 106, 634a.

Stevens, T.J. and Arkin, I.T. (2000) Do more complex organisms have a greater proportion of membrane proteins in their genomes? Proteins Struct. Funct. Genet., 39, 417–420.

Tusnady, G.E. et al. (2004) Transmembrane proteins in the Protein Data Bank: identification and classification. Bioinformatics, 20, 2964–2972.

Valas, R.E. et al. (2009) Nothing about protein structure classification makes sense except in the light of evolution. Curr. Opin. Struct. Biol., 19, 329–334.

Wetlaufer, D.B. (1973) Nucleation, rapid folding, and globular intrachain regions in proteins. Proc. Natl. Acad. Sci., 70, 697–701.

White, S.H. (2009) Biophysical dissection of membrane proteins. Nature, 459, 344–346.

Wu, L.C. et al. (2008) Autonomous subdomains in protein folding. Protein Sci., 3, 369–371.

Xu, J. and Zhang, Y. (2010) How significant is a protein structure similarity with TM-score = 0.5? Bioinformatics, 26, 889–895.

Zhang, Y. and Skolnick, J. (2004) Scoring function for automated assessment of protein structure template quality. Proteins Struct. Funct. Genet., 57, 702–710.

